# Lack of evidence for cross-frequency phase-phase coupling between theta and gamma oscillations in the hippocampus

**DOI:** 10.1101/045963

**Authors:** Robson Scheffer-Teixeira, Adriano BL Tort

**Affiliations:** Brain Institute, Federal University of Rio Grande do Norte, Natal, RN 59056-450, Brazil

## Abstract

Phase-amplitude coupling between theta and multiple gamma sub-bands hallmarks hippocampal activity and is believed to take part in information routing. More recently, theta and gamma oscillations were also reported to exhibit reliable phase-phase coupling, or n:m phase-locking. The existence of n:m phase-locking suggests an important mechanism of neuronal coding that has long received theoretical support. However, here we show that n:m phase-locking (1) is much lower than previously reported, (2) highly depends on epoch length, (3) does not statistically differ from chance (when employing proper surrogate methods), and that (4) filtered white noise has similar n:m scores as actual data. Moreover, (5) the diagonal stripes in theta-gamma phase-phase histograms of actual data can be explained by theta harmonics. These results point to lack of theta-gamma phase-phase coupling in the hippocampus, and suggest that studies investigating n:m phase-locking should rely on appropriate statistical controls, otherwise they could easily fall into analysis pitfalls.

## Introduction

Local field potentials (LFPs) exhibit oscillations of different frequencies, which may co-occur and also interact with one another (Jensen and Colgin, 2007; Tort et al., 2010; Hyafil et al., 2015). Cross-frequency phase-amplitude coupling between theta and gamma oscillations has been well described in the hippocampus, whereby the instantaneous amplitude of gamma oscillations depends on the instantaneous phase of theta (Scheffer-Teixeira et al., 2012; Schomburg et al., 2014). More recently, Belluscio et al. (2012) reported that theta and gamma oscillations also exhibit reliable phase-phase coupling, in which multiple gamma cycles are consistently entrained within one cycle of theta. The existence of multiple types of cross-frequency coupling suggests that the brain may use different coding strategies to transfer multiplexed information.

Coherent oscillations are believed to take part in network communication by allowing opportunity windows for the exchange of information (Varela et al., 2001; Fries, 2005). Standard phase coherence measures the constancy of the phase difference between two oscillations of the same frequency (Lachaux et al., 1999; Hurtado et al., 2004), and it has been associated with cognitive processes such as decision-making (DeCoteau et al., 2007; Montgomery and Buzsáki, 2007; Nácher et al., 2013). Similarly to coherence, cross-frequency phase-phase coupling, or n:m phase-locking, also relies on assessing the constancy of the difference between two phase time series (Tass et al., 1998). However, in this case the original phase time series are accelerated, so that their instantaneous frequency can match. Formally, n:m phase-locking occurs when Δ*φ*_*nm*_(*t*) = *n* * *φ*_*B*_(*t*) – *m* * *φ*_*A*_(*t*) is constant, where *n* * *φ*_*B*_ (*m* * *φ*_*A*_) denotes the phase of oscillation B (A) accelerated n (m) times (Tass et al., 1998). For example, the instantaneous phase of theta oscillations at 8 Hz needs to be accelerated 5 times to match in frequency a 40-Hz gamma. A 1:5 phase-phase coupling is then said to occur if the instantaneous phase of theta accelerated 5 times has constant difference to the instantaneous gamma phase; or, in other words, if 5 gamma cycles have fixed phase relationship to 1 theta cycle.

Cross-frequency phase-phase coupling has previously been hypothesized to take part in memory processes (Lisman and Idiart, 1995; Jensen and Lisman, 2005; Lisman, 2005; Schack and Weiss, 2005; Sauseng et al., 2008; Sauseng et al., 2009; Holtz et al., 2010; Fell and Axmacher, 2011). The recent findings of Belluscio et al. (2012) are important because they suggest that the hippocampus indeed uses such a mechanism. However, in the present work we question the existence of theta-gamma phase-phase coupling. By analyzing simulated and actual hippocampal LFPs, we show that the levels of n:m phase-locking between theta and gamma oscillations are not greater than chance.

## Results

We first certified that we could reliably detect n:m phase-locking when present. To that end, we simulated a 20-s LFP signal containing white noise, high-amplitude theta (8 Hz) and low-amplitude gamma (40 Hz) oscillations (Figure 1A, top). The phase time series for theta and gamma were obtained from the simulated LFP after band-pass filtering (Figure 1A, bottom). Figure 1B depicts three versions of accelerated theta phases (m=3, 5 and 7) along with the instantaneous gamma phase (n=1). Also shown is the time series of the difference between gamma and accelerated theta phases (Δ*φ*_*nm*_). The instantaneous phase difference is roughly constant only for m=5; when m=3 or 7, it changes over time, precessing forwards (m=3) or backwards (m=7) at a rate of 16 Hz. Consequently, Δ*φ*_*nm*_ distribution is uniform over 0 and 2π for m=3 or 7, but highly concentrated for m=5 (Figure 1C). The concentration (or “constancy”) of the phase difference distribution is commonly used as a metric of n:m phase-locking. This metric is defined as the length of the mean resultant vector (R_n:m_) over unitary vectors whose angle is the instantaneous phase difference (*e*^*i*Δ*φ*_*nm*_(*t*)^), and thereby it varies between 0 and 1. For any pair of filtered LFPs, an R_n:m_ “curve” can be obtained by varying m for n=1 fixed. When filtered at theta and gamma, the simulated LFP exhibited a prominent peak for n:m = 1:5 (Figure 1D, left), which shows that R_n:m_ successfully detects n:m phase-locking. Importantly, in this first case gamma frequency was an integer multiple of theta frequency (5 × 8 Hz = 1 × 40 Hz), or, in other words, gamma frequency was a harmonic of theta frequency. We next repeated the analysis above for a similar LFP, but simulated with a slightly different gamma frequency, set to 39.9 Hz. In this second case the R_n:m_ curve exhibited no prominent peak (Figure 1D, right). Therefore, R_n:m_ is highly sensitive to the exact peak frequency of the analyzed oscillations. This fact alone casts some doubt on the existence of n:m phase-locking between actual brain oscillations, as their peak frequencies are unlikely to be perfect integer multiples.

**Figure 1.**
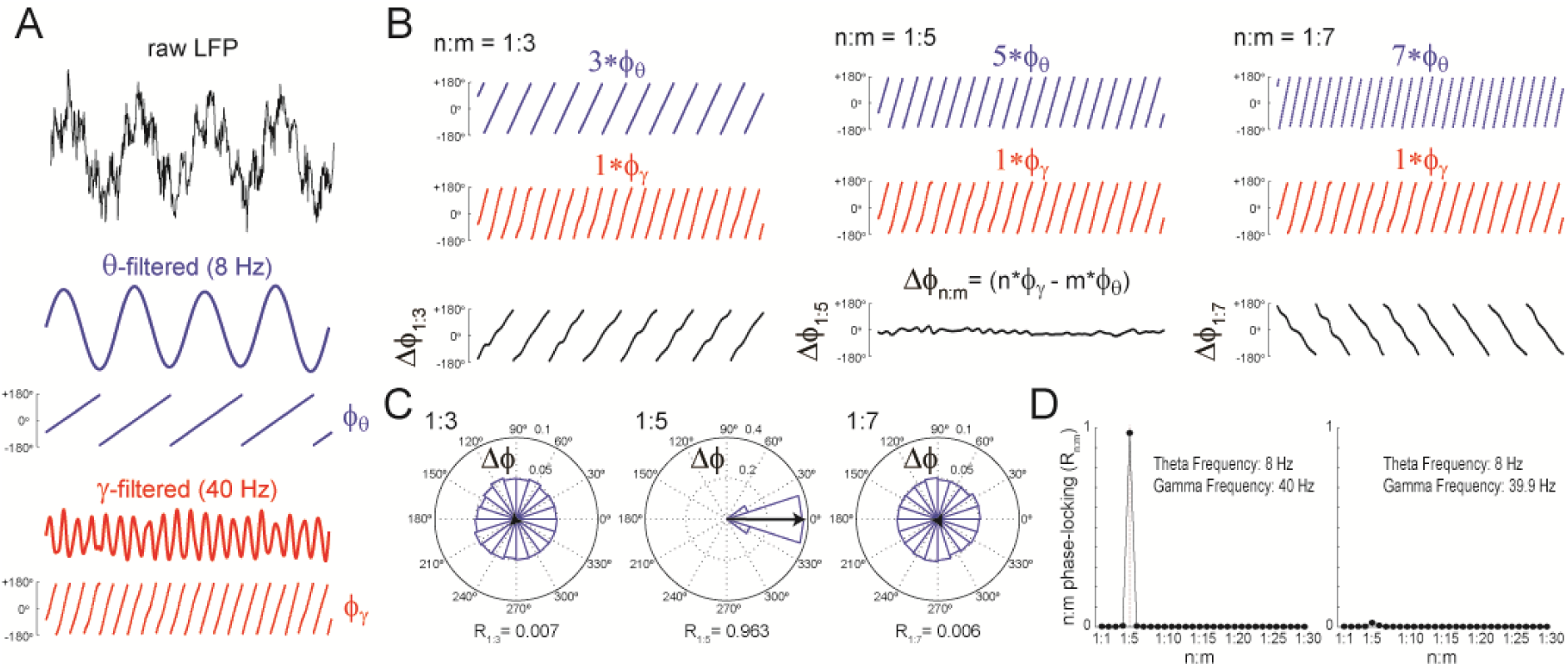
Measuring cross-frequency phase-phase coupling. (A) Top trace shows 500 ms of simulated LFP containing white noise, theta (8 Hz) and gamma (40 Hz) oscillations. Bottom traces show theta-(blue) and gamma-(red) filtered signals and their instantaneous phases (filter bands: 6-10 Hz for theta and 38-42 Hz for gamma). (B) Top blue traces show the instantaneous theta phase for the same period as in A but accelerated *m* times, where m= 3 (left), 5 (middle) and 7 (right). Middle red traces reproduce the instantaneous gamma phase (i.e., n=1). Bottom black traces show the instantaneous phase difference between gamma and accelerated theta phases (Δ*φ*_*nm*_). Notice roughly constant Δ*φ*_*nm*_ only when theta is accelerated m=5 times, which indicates 1:5 phase-locking. (C)Δ*φ*_*nm*_ distributions. Notice uniform distributions for n:m=1:3 and 1:7, and a highly concentrated distribution for n:m=1:5. The black arrow represents the mean resultant vector for each case (see Material and Methods). The length of this vector (R^n:m^) measures the level of n:m phase-locking. (D)Left, phase-locking levels for a range n:m ratios for the signal in A. Right, same as before but for a simulated LFP in which gamma frequency was set to 39.9 Hz.

We next sought to understand why previous studies found theta-gamma phase-phase coupling in actual LFP recordings despite the observations above (Belluscio et al., 2012; Zheng et al., 2013; Xu et al., 2013; Stujenske et al., 2014; Xu et al., 2015; Zheng et al., 2016). We started by analyzing white-noise signals, in which by definition there is no structured activity; in particular, the spectrum is flat and there is no n:m phase-locking. R_n:m_ values measured from white noise should be regarded as chance levels. We band-pass filtered white-noise signals to extract the instantaneous phase of theta (θ: 4 – 12 Hz) and of multiple gamma bands (Figure 2A): slow gamma (γs: 30 – 50 Hz), middle gamma (γm: 50 – 90 Hz), and fast gamma (γf: 90 – 150 Hz). For each θ–γpair, we constructed n:m phase-locking curves for epochs of 1 and 10 seconds, with n=1 fixed and m varying from 1 to 25 (Figure 2B). In each case, phase-phase coupling was high within the ratio of the analyzed frequency ranges: R_n:m_ peaked at m=4-6 for θ–γs, at m=7-11 for θ–γM, and at m=12-20 for θ–γ_F_. Therefore, the existence of a “bump” in the n:m phase-locking curve may merely reflect the ratio of the filtered bands and should not be considered as evidence for cross-frequency phase-phase coupling; even filtered white-noise signals exhibit such a pattern.

**Figure 2.**
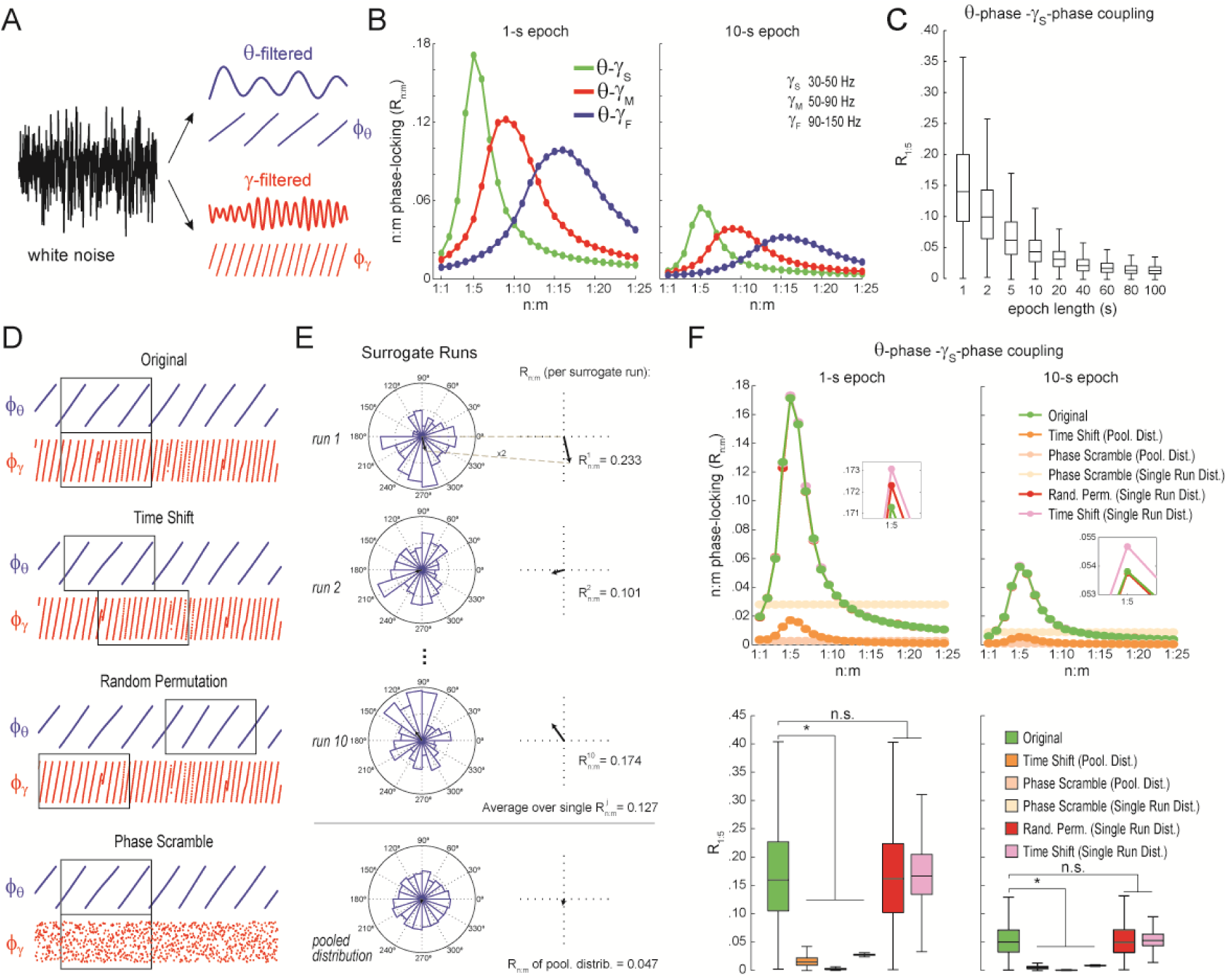
Detection of spurious n:m phase-locking in white-noise signals due to inappropriate surrogate-based statistical testing. (A)Example white-noise signal (black) along with its theta-(blue) and gamma-(red) filtered components. The corresponding instantaneous phases are also shown. (B)n:m phase-locking levels for 1-(left) and 10-second (right) epochs, computed for noise filtered at theta (θ; 4 – 12 Hz) and at three gamma bands: slow gamma (γs; 30 – 50 Hz), middle gamma (γM; 50 – 90 Hz) and fast gamma (γF; 90 – 150 Hz). Notice R_n:m_ peaks in each case. (C)Boxplot distributions of θ-γs R1:5 values for different epoch lengths (n=2100 simulations per epoch length). (D)Overview of surrogate techniques. See text for details. (E)Top panels show representative Δ*φ*_*nm*_ distributions for single surrogate runs (*Time Shift*; 10 runs of 1-s epochs), along with the corresponding R_n:m_ values. The bottom panel shows the pooled Δ*φ*_*nm*_ distribution; the R_n:m_ of the pooled distribution is lower than the R_n:m_ of single runs (compare with values for 1‐ and 10-s epochs in panel C). (F)Top, n:m phase-locking levels computed for 1-(left) or 10-s (right) epochs using either the *Original* or 5 surrogate methods (insets are a zoomed view of R_n:m_ peaks). Bottom, R1:5 values for white noise filtered at 9 and γs. Original R_n:m_ values are not different from R_n:m_ values obtained from single surrogate runs of *Random Permutation* and *Time Shift* procedures. Less conservative surrogate techniques provide lower R_n:m_ values and lead to the spurious detection of θ-γs phase-phase coupling in white noise. ^*^<0.01, n=2100 per distribution, one-way ANOVA with Bonferroni post-hoc test.

Qualitatively similar results were found for 1‐ and 10-s epochs; however, R_n:m_ values were considerably lower for the latter (Figure 2B). In fact, for any fixed n:m ratio and frequency pair, R_n:m_ decreased as a function of epoch length (see Figure 2C for θ–γ_S_ and R_1:5_): the longer the white-noise epoch the more the phase difference distribution becomes uniform. In other words, as standard phase coherence (Vinck et al., 2010) and phase-amplitude coupling (Tort et al., 2010), phase-phase coupling also has positive bias for shorter epochs; an adequate time length is required to average out noise and reveal true coupling, if any. As a corollary, notice that false-positive coupling may be detected if control (surrogate) epochs are longer than the original epoch.

We next investigated the reliability of surrogate methods for detecting n:m phase-locking (Figure 2D). The *“Original”* R_n:m_ value uses the same time window for extracting theta and gamma phases (Figure 2D, upper panel). Belluscio et al. (2012) employed a *“Time Shift”* procedure for creating surrogate epochs, in which the time window for gamma phase is randomly shifted between 1 to 200 ms from the time window for theta phase (Figure 2D, upper middle panel). A variant of this procedure is the *“Random Permutation”*, in which the time window for gamma phase is randomly chosen (Figure 2D, lower middle panel). Finally, in the *“Phase Scramble”* procedure, the timestamps of the gamma phase time series are randomly shuffled (Figure 2D, lower panel); clearly, the latter is the least conservative. For each surrogate procedure, R_n:m_ values were obtained by two approaches: *“Single Run”* and *“Pooled”* (Figure 2E). In the first approach, each surrogate run (e.g., a time shift or a random selection of time windows) produces one R_n:m_ value (Figure 2E, top panels). In the second, Δ*φ*_*nm*_ from several surrogate runs are first pooled, then a single R_n:m_ value is computed from the pooled distribution (Figure 2E, bottom panel). As illustrated in Figure 2E, R_n:m_ computed from a pool of surrogate runs is much smaller than when computed for each individual run. This is due to the dependence of R_n:m_ on the epoch length: pooling instantaneous phase differences across 10 runs of 1-s surrogate epochs is equivalent to analyzing a single surrogate epoch of 10 seconds. And the longer the analyzed epoch, the more the noise is averaged out and the lower the R_n:m_. Therefore, pooled surrogate epochs summing up to 10-s of total data have lower R_n:m_ than any individual 1-s surrogate epoch.

No phase-phase coupling should be detected in white noise, and therefore *Original* R_n:m_ values should not differ from chance. However, as shown in Figure 2F for θ–γs as an illustrative case (similar results hold for any frequency pair), θ–γs phase-phase coupling in white noise is considered statistically significant when compared to phase-scrambled surrogates (for either *Single Run* or *Pooled* distributions). This was true for surrogate epochs of any length, although the longer the epoch, the lower the actual and the surrogate R_n:m_ values, as expected (compare right and left panels of Figure 2F). *Pooled* R1:5 distributions derived from either time-shifted (Figure 2F) or randomly permutated epochs (not shown) also led to the detection of false positive θ–γs phase-phase coupling. On the other hand, *Original* R_n:m_ values were not statistically different from chance distributions when these were constructed from *Single Run* R_n:m_ values for either *Time Shift* and *Random Permutation* surrogate procedures (Figure 2F). We conclude that neither scrambling phases nor pooling individual surrogate epochs should be employed for statistically evaluating n:m phase-locking. Chance distributions should be derived from surrogate epochs of the same length as the original epoch and which preserve phase continuity.

We next proceeded to analyze hippocampal CA1 recordings from 7 rats, focusing on periods of prominent theta activity (active waking and REM sleep). We found strikingly similar results between white noise and actual LFP data. Namely, R_n:m_ curves peaked at n:m ratios according to the filtered bands, and R_n:m_ values were lower for longer epochs (Figure 3A; compare with Figure 2B). We also computed phase-phase plots (2D histograms of theta phase vs gamma phase) and observed diagonal stripes as reported in Belluscio et al. (2012) (see also Zheng et al., 2016), which seem to suggest phase-phase coupling (Figure 3B). However, as shown in Figure 3C, the original R_n:m_ values were not statistically different from a proper surrogate distribution (*Random Permutation/Single Run*). Noteworthy, as with white-noise data, false positive phase-phase coupling would be inferred if an inadequate surrogate method were employed (*Time Shift/Pooled*) (Figure 3C).

**Figure 3.**
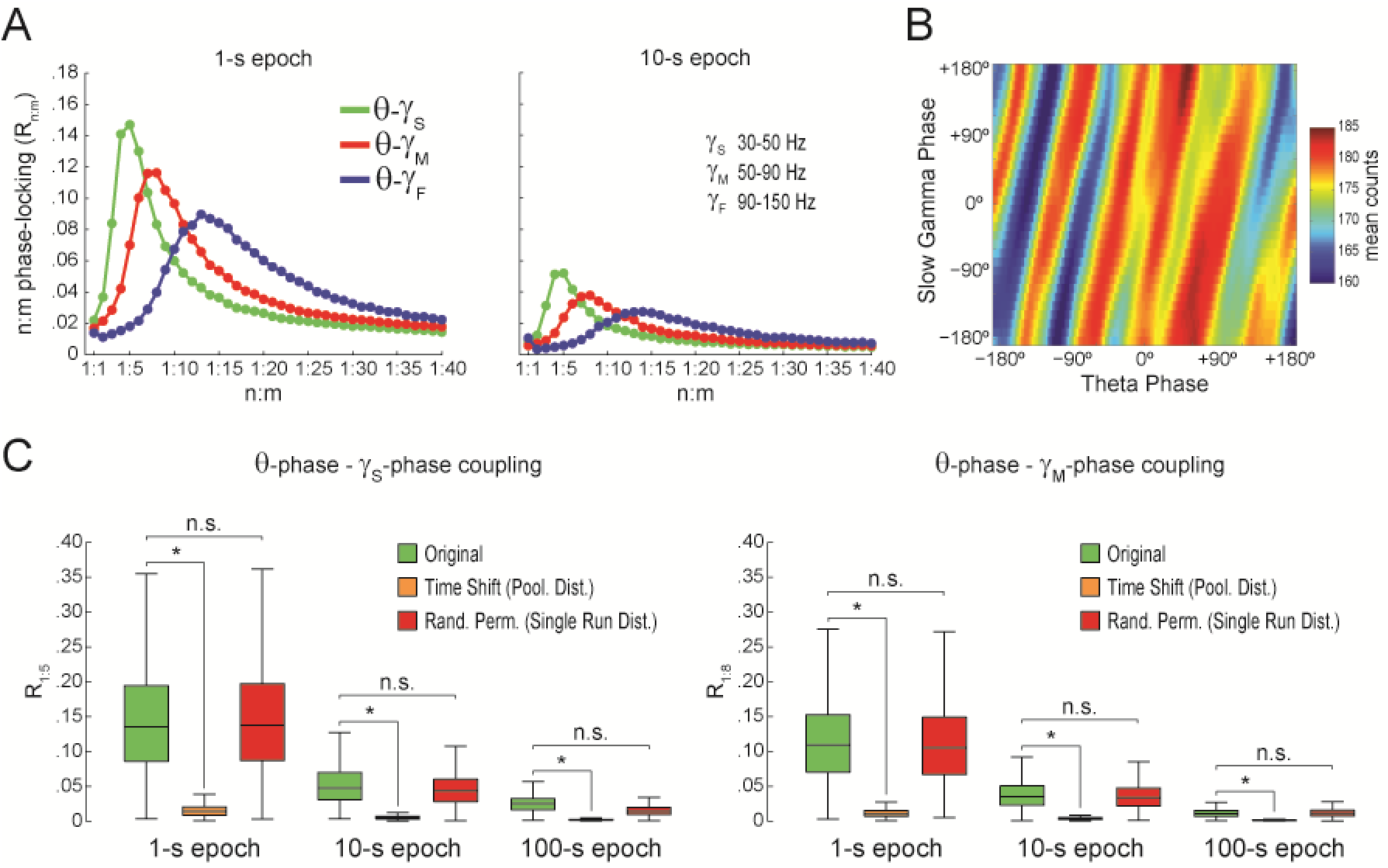
Spurious detection of theta-gamma phase-phase coupling in the hippocampus. (A) n:m phase-locking levels for actual hippocampal LFPs. Compare with Figure 2B. (B) Phase-phase plot for theta and slow gamma (average over animals). Notice diagonal stripes suggesting phase-phase coupling. (C) Original and surrogate distributions of R_n:m_ values for slow (R_1:5_;left) and middle gamma (R_1:8_; right) for different epoch lengths. The original data is significantly higher than the pooled surrogate distribution, but indistinguishable from the distribution of surrogate values computed using single runs. Similar results hold for fast gamma. ^*^p<0.01, n=7 animals, Friedman’s test with Nemenyi post-hoc test.

Based on the above, we concluded that there is no evidence for n:m phase-locking in actual hippocampal LFPs. We next sought to investigate what causes the diagonal stripes in phase-phase plots that are appealingly suggestive of 1:5 coupling (Figure 3B). In Figure 4 we analyze a representative LFP with prominent theta oscillations at ~7 Hz recorded during REM sleep. We constructed phase-phase plots using LFP components narrowly filtered at theta and its harmonics: 14, 21, 28 and 35 Hz. For each harmonic frequency, the phase-phase plot exhibited diagonal stripes whose number was determined by the harmonic order (i.e., the 1^st^ harmonic exhibited two stripes, the 2^nd^ harmonic three stripes, the 3^rd^, four stripes and the 4^th^, five stripes; Figure 4B i-iv). Interestingly, when the LFP was filtered at a broad gamma band (30-90 Hz), we observed 5 diagonal stripes, the same number as when narrowly filtering at 35 Hz; moreover, both gamma and 35-Hz filtered signals exhibited the exact same phase lag (Figure 4B iv-v). Therefore, these results indicate that the diagonal stripes in phase-phase plots are due to theta harmonics and not to genuine gamma activity. Signals filtered at the gamma band are likely to exhibit as many stripes as expected for the first theta harmonic falling within the filtered band (Figure 5). Similar results were observed in all animals during both active awaking and REM states. Importantly, as in Belluscio et al. (2012), phase-phase plots constructed using data from multiple time-shifted epochs exhibited no diagonal stripes (Figure 4B vi). This is because different time shifts lead to different phase lags; the diagonal stripes of individual surrogate runs that could otherwise be apparent cancel each other out when combining data across multiple runs of different lags (Figure S1).

**Figure 4.**
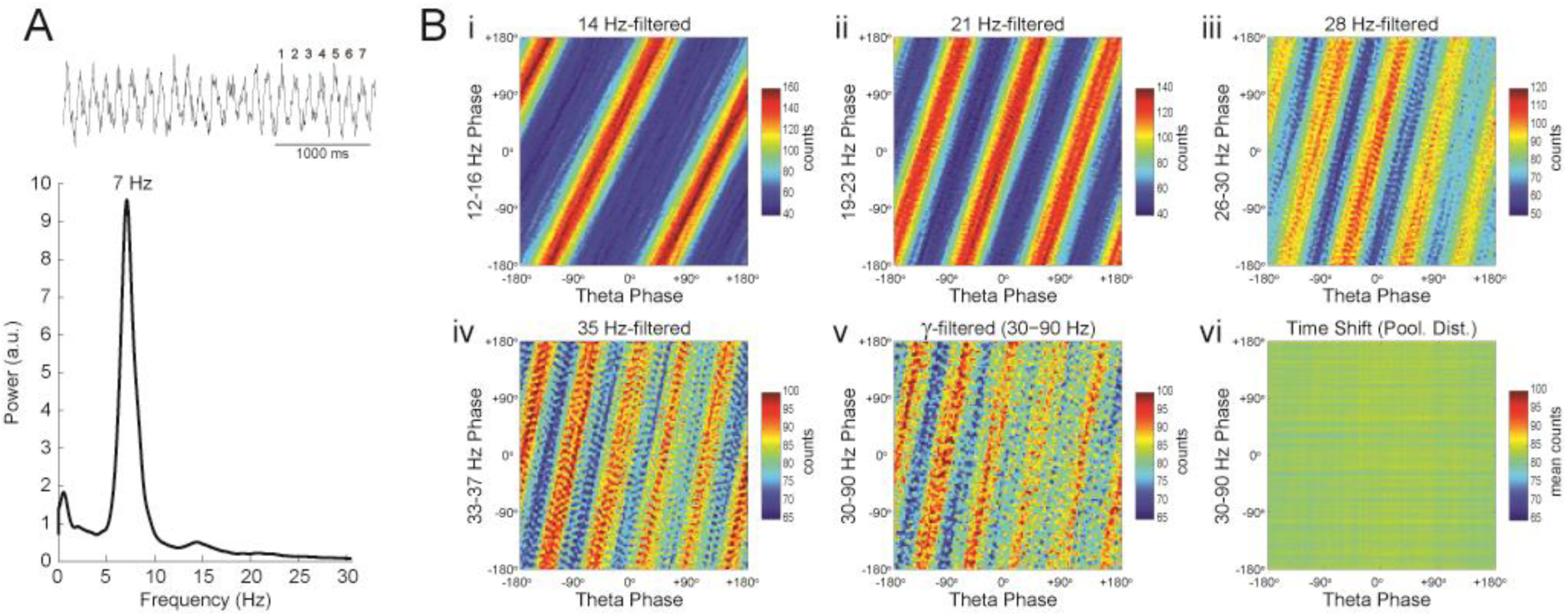
Phase-phase coupling between theta and gamma oscillations is confounded by theta harmonics. (A) Top, representative LFP epoch exhibiting prominent theta activity (~7 Hz) during REM sleep. Bottom, power spectral density. (B) Phase-phase plots for theta and LFP band-pass filtered at harmonic frequencies (14, 21, 28 and 35 Hz), computed using 20 minutes of concatenated REM sleep. Also shown are phase-phase plots for the conventional gamma band (30 – 90 Hz) and for pooled surrogate runs. Notice that the former mirrors the phase-phase plot of the 4^th^ theta harmonics (35 Hz).

**Figure 5.**
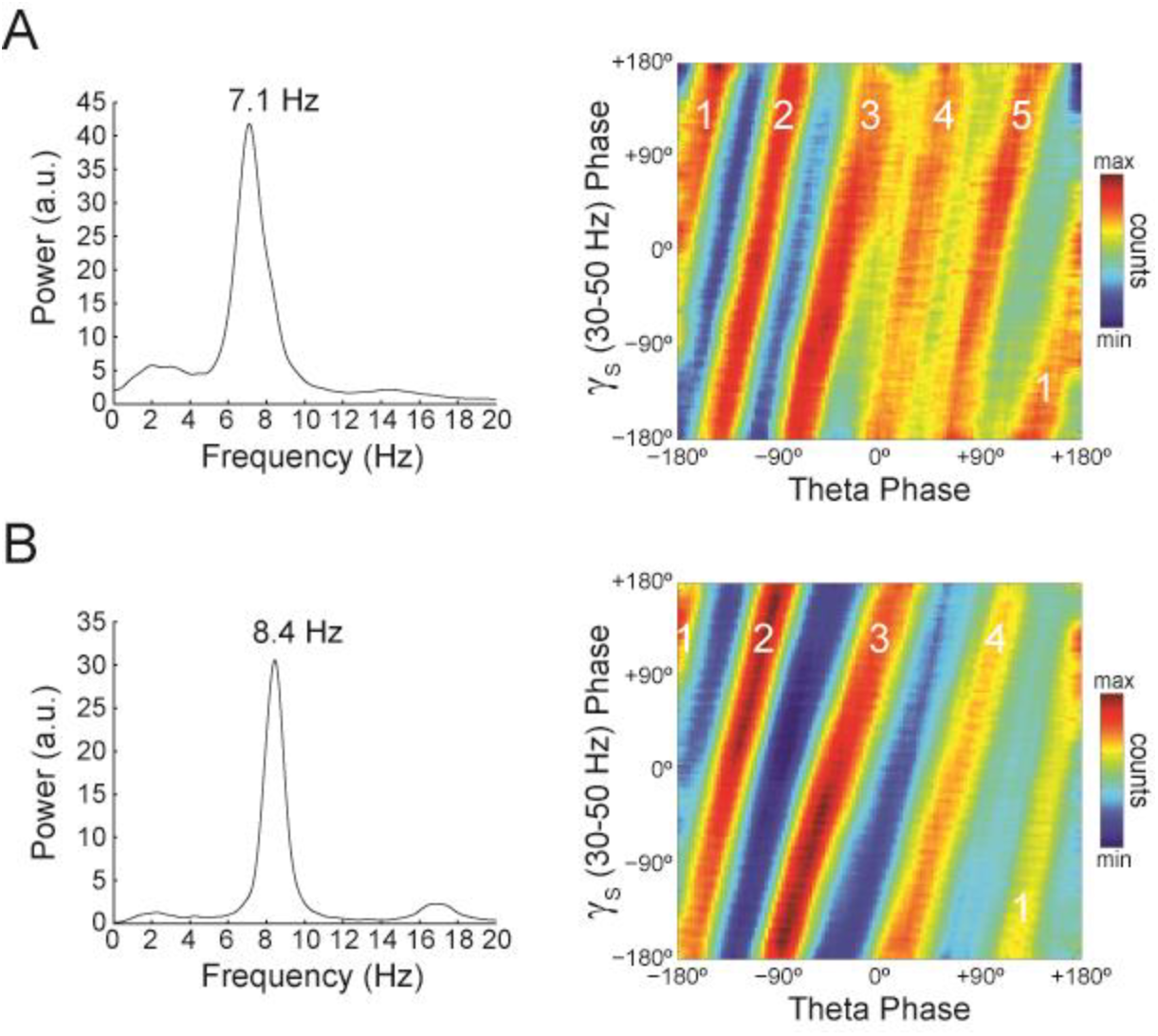
The number of stripes in phase-phase plots is determined by the frequency of the first theta harmonic within the filtered gamma range. (A)Representative example in which theta has peak frequency of 7.1 Hz. The phase-phase plot between theta and slow gamma (30-50 Hz) exhibits 5 stripes, since the 4^th^ theta harmonic (35.5 Hz) is the first to fall within 30 and 50 Hz. (B)Example in which theta has peak frequency of 8.4 Hz and the phase-phase plot exhibits 4 stripes, which correspond to the 3^rd^ theta harmonic (33.6 Hz).

Our findings also held true when employing a pair-wise phase consistency metric (Vinck et al., 2010) (not shown), or when estimating theta phase through a method based on interpolating phase values between 4 points of the theta cycle (trough, ascending, peak and descending points) as performed in Belluscio et al. (2012) (Figure S2). The latter was somewhat expected since the phase-phase coupling results in Belluscio et al. (2012) did not depend on this particular method of phase estimation (see their Figure 6Ce). We further confirmed our results by analyzing data from 3 additional rats recorded in an independent laboratory (Figure S3; see Material and Methods). In addition, we also found no theta-gamma phase-phase coupling neither in LFPs from other hippocampal layers than s. *pyramidale* (Figure S4), nor in neocortical LFPs (not shown), nor in current-source density (CSD) signals (Figure S4), nor in independent components that isolate activity of specific gamma sub-bands (Schomburg et al., 2014) (Figure S5), nor in transient gamma bursts (Figure S6).

In all, our results point to lack of evidence for cross-frequency phase-phase coupling in the rat hippocampus, and further show that the diagonal stripes observed in phase-phase plots previously attributed to theta-gamma coupling are actually explained by theta harmonics alone.

## Discussion

Theta and gamma oscillations hallmark hippocampal activity during active exploration and REM sleep (Buzsáki et al., 2003; Csicsvari et al., 2003; Montgomery et al., 2008). Theta and gamma are well known to interact by means of phase-amplitude coupling, in which the instantaneous gamma amplitude waxes and wanes as a function of theta phase (Bragin et al., 1995; Scheffer-Teixeira et al., 2012; Caixeta et al., 2013). This particular type of cross-frequency coupling has been receiving large attention and related to functional roles (Canolty and Knight, 2010;Hyafil et al., 2015). In addition to phase-amplitude coupling, theta and gamma oscillations can potentially interact in many other ways (Jensen and Colgin, 2007; Hyafil et al., 2015). For example, the power of slow gamma oscillations may be inversely related to theta power (Tort et al., 2008), suggesting amplitude-amplitude coupling. Recently, Belluscio et al. (2012) reported that theta and gamma would also couple by means of n:m phase-locking. Among other implications, this finding was taken as evidence for network models of working memory (Lisman and Idiart, 1995; Jensen and Lisman, 2005; Lisman, 2005) and for a role of basket cells in generating cross-frequency coupling (Belluscio et al., 2012; Buzsáki and Wang, 2012). However, our results show that hippocampal levels of n:m phase-locking do not differ from chance, and further suggest that previous work may have spuriously detected phase-phase coupling due to an improper use of surrogate methods.

### Statistical inference of phase-phase coupling

When attempting to reproduce Belluscio et al. (2012), we noticed that our R_n:m_ values were much smaller than reported in their study. Namely, for each individual animal, R_n:m_ was lower than 0.02 when analyzing all epochs of active waking or REM sleep. We suspected that this could be due to differences in the duration of the analyzed epochs between both studies. We then investigated the dependence of R_n:m_ on epoch length, and found a strong positive bias for shorter epochs. In addition, we found that R_n:m_ values exhibit greater variability as epoch length decreases for both white noise and actual data. Since theta and gamma peak frequencies are not constant in these signals, the longer the epoch, the more the theta and gamma peak frequencies are allowed to fluctuate and the more apparent the lack of coupling. On the other hand, Δ*φ*_*nm*_ distribution becomes less uniform for shorter epochs. The dependence of n:m phase-coupling metrics on epoch length has important implications in designing surrogate epochs for testing the statistical significance of actual R_n:m_ values. As demonstrated here, spurious detection of phase-phase coupling may occur if surrogate epochs are longer than the original epoch. This is clearly the case when one lumps together several surrogate epochs before computing R_n:m_. When employing proper controls, our results show that R_n:m_ values of real data do not differ from surrogate values, and are actually similar to R_n:m_ values obtained for white noise (irrespective of the epoch length; compare Figures 2 and 3).

Belluscio et al. (2012) have not tested the significance of individual R_n:m_ values; instead, they statistically inferred the existence of n:m phase-locking by comparing empirical phase-phase plots with those obtained from the average of 1000 time-shifted surrogate runs (they established a significance threshold for each phase-phase bin based on the mean and standard deviation of surrogate counts in that bin). Belluscio et al. (2012) assumed that the diagonal stripes observed in phase-phase plots were due to theta-gamma coupling; no such stripes were apparent in phase-phase plots constructed from the average over surrogate epochs (c.f. their Figure 6). However, here we show that the presence of diagonal stripes in phase-phase plots is not sufficient to conclude the existence of phase-phase coupling; actually, these stripes can be explained by theta harmonics alone. Accordingly, the analysis of synthetic LFPs having no gamma oscillations and theta simulated as a saw-tooth wave leads to similar results (Figure S7). That is, even though by definition the theta saw-tooth wave has no theta-gamma phase-phase coupling, filtering at the gamma band will reflect the harmonic of theta with largest amplitude within the filtered band and give rise to diagonal stripes in phase-phase plots. These would be spuriously deemed as statistically significant if we were to use the same statistical analysis as in Belluscio et al. (2012), since phase-phase plots obtained from the average of 1000 surrogates with shuffled phase lags exhibit no diagonal stripes (Figure S7).

### Lack of evidence vs evidence of non-existence

One could argue that we were unable to detect statistically significant theta-gamma phase-phase coupling because we did not analyze a proper dataset, or else because phase-phase coupling would only occur during certain behavioral states not investigated here. We disagree with these arguments for the following reasons: (1) we could reproduce our results using data from the same laboratory as Belluscio et al. (2012) (Figure S3), and (2) we examined the exact same behavioral states as in their study (active waking and REM sleep). One could also argue that there exists multiple gammas, and that different gamma types are most prominent in different hippocampal layers (Colgin et al., 2009; Scheffer-Teixeira et al., 2012; Tort et al., 2013; Schomburg et al., 2014; Lasztóczi and Klausberger, 2014); therefore, theta-gamma phase-phase coupling could exist in other hippocampal layers not investigated here. We also disagree with this possibility because: (1) we examined the same hippocampal layer in which theta-gamma phase-phase coupling was reported to occur (Belluscio et al., 2012); (2) we have also analyzed other layers in addition to s. *pyramidale* (we recorded LFPs using 16-channel silicon probes, see Material and Methods) and found lack of phase-phase coupling in all hippocampal layers (Figure S4); (3) we have actually also found lack of theta-gamma phase-phase coupling in parietal and entorhinal cortex recordings (not shown). Moreover, similar results hold when (4) filtering LFPs within any gamma sub-band (Figure 3 and Figures S2 to S6), (5) analyzing CSD signals (Figure S4), or (6) analyzing independent components that maximize activity within particular gamma sub-bands (Schomburg et al., 2014) (Figure S5). Finally, one could argue that gamma oscillations are not continuous but transient, and that assessing phase-phase coupling between theta and transient gamma bursts would require a different type of analysis than employed here. Regarding this argument, we once again stress that we used the exact same methodology as originally used to detect theta-gamma phase-phase coupling (Belluscio et al., 2012). Nevertheless, we also ran analysis only taking into account periods in which gamma amplitude was >2SD above the mean (“gamma bursts”) and found no statistically significant phase-phase coupling (Figure S6).

As showcased in Figure 1D, n:m phase-locking is highly sensitive to the exact peak frequencies of the analyzed oscillations. In those examples, there is no longer 1:5 coupling if gamma peak frequency differs by 0.1 Hz from 40 Hz (that is, n^*^γ*freq* = 1^*^39.9 Hz differs 0.1 Hz from m^*^θ*freq* = 5^*^8 Hz). This high R_n:m_ sensitivity reflects a mathematical property of sinusoidal basis functions: sine of different frequencies are orthogonal, irrespective of the magnitude of the frequency difference. However, we note that n:m phase-locking would still be detected in this example if we had used n=80 and m=399, or, alternatively, if we had used n=1 and m=39.9/8 = 4.9875. However, we restricted the analysis R_n:m_ to low integer values since the exact same was done in previous studies (Belluscio et al., 2012; Zheng et al., 2013; Xu et al., 2013; Stujenske et al., 2014; Xu et al., 2015; Zheng et al., 2016). In hindsight, however, it would have been better if the previous studies had not used integer values of m (for n=1 fixed) in order to avoid contamination with harmonics. At any event, we also deem unlikely that two genuine brain oscillations would differ in peak frequency by a perfect rational number.

The high sensitivity of R_n:m_ on exact peak frequencies casts some doubt as to whether theta-gamma n:m phase-locking can ever exist in the brain. Contrary to this intuition, however, following Belluscio et al. (2012) other studies also reported theta-gamma phase-phase coupling in the rodent hippocampus (Zheng et al., 2013; Xu et al., 2013; Xu et al., 2015; Zheng et al., 2016) and amygdala (Stujenske et al., 2014). In addition, human studies had previously reported theta-gamma phase-phase coupling in scalp EEG (Sauseng et al., 2008; Sauseng et al., 2009; Holtz et al., 2010). Nonetheless, most of these studies have not tested the statistical significance of coupling levels against chance (Sauseng et al., 2008; Sauseng et al., 2009; Holtz et al., 2010; Zheng et al., 2013; Xu et al., 2013; Xu et al., 2015; Stujenske et al., 2014), while Zheng et al. (2016) used similar statistical inferences as in Belluscio et al. (2012). We further note that epoch length was often not informed in the animal studies.

Since it is philosophically impossible to prove absence of an effect, the burden of proof should be placed on demonstrating that a true effect exists. In this sense, and to the best of our knowledge, none of previous research investigating theta-gamma phase-phase coupling has properly tested R_n:m_ against chance. Many studies have focused on comparing changes in n:m phase-locking levels, but we believe these can be influenced by other variables such as changes in power. Interestingly, in their pioneer work, Tass and colleagues used filtered white noise to construct surrogate distributions and did not find significant n:m phase-locking among brain oscillations (Tass et al., 1998; Tass et al., 2003).

### Implications for models of neural coding by theta-gamma coupling

In 1995, Lisman and Idiart proposed an influential model in which theta and gamma oscillations would interact to produce a neural code. The theta-gamma coding model has since been improved (Jensen and Lisman, 2005; Lisman, 2005; Lisman and Buzsáki, 2008), but its essence remains the same (Lisman and Jensen, 2013): nested gamma cycles would constitute memory slots, which are parsed at each theta cycle. Accordingly, Lisman and Idiart (1995) hypothesized that working memory capacity (7±2) is determined by the number of gamma cycles per theta cycle.

Both phase-amplitude and phase-phase coupling between theta and gamma have been considered experimental evidence for such coding scheme (Lisman and Buzsáki, 2008; Sauseng et al., 2009; Axmacher et al., 2010; Belluscio et al., 2012; Lisman and Jensen, 2013; Hyafil et al., 2015; Rajji et al., 2016). In the case of phase-amplitude coupling, the modulation of gamma amplitude within theta cycles would instruct a reader network when the string of items represented in different gamma cycles starts and terminates. On the other hand, the precise ordering of gamma cycles within theta cycles that is consistent across theta cycles would imply phase-phase coupling; indeed, n:m phase-locking is a main feature of computational models of sequence coding by theta-gamma coupling (Lisman and Idiart, 1995; Jensen and Lisman, 1996; Jensen et al., 1996). In contrast to these models, however, the absence of theta-gamma phase-phase coupling reported here shows that the theta phases in which gamma cycles begin/end are not fixed across theta cycles, which is to say that gamma cycles are not precisely timed but rather drift; in other words, gamma is not a clock (Burns et al., 2011).

If theta-gamma neural coding exists, our results suggest that the precise location of gamma memory slots within a theta cycle is not required for such a code, and that the ordering of the represented items would be more important than the exact spike timing of the cell assemblies that represent the items (Lisman and Jensen, 2013).

## Conclusion

In summary, while absence of evidence is not evidence of absence, our results strongly suggest that theta-gamma phase-phase coupling does not exist in the hippocampus. We believe that evidence in favor of n:m phase locking in other brain regions and signals should be revisited and, whenever suitable, checked against the more conservative surrogate techniques outlined here.

## Material and Methods

*Animals and surgery.* All procedures were approved by our local institutional ethics committee and were in accordance with the National Institutes of Health guidelines. We used 7 male Wistar rats (2–3 months; 300-400 g) from our breeding colony, kept under 12h/12h dark-light cycle. We recorded from the dorsal hippocampus through either multi-site linear probes (n=6 animals; 4 probes had 16 4320–μm^2^ contacts spaced by 100 μm; 1 probe had 16 703–μm^2^ contacts spaced by 100 μm; 1 probe had 16 177–μm^2^ contacts spaced by 50 μm; all probes from NeuroNexus) or single wires (n=1 animal; 50–μm diameter) inserted at AP -3.6 mm and ML 2.5 mm. Results shown in the main figures were obtained for LFP recordings from the CA1 pyramidal cell layer, identified by depth coordinate and characteristic electrophysiological benchmarks such as highest ripple power (see Figure S4 for an example). Similar results were obtained for recordings from other hippocampal layers (Figure S4).

We also analyzed data from 3 additional rats downloaded from the Collaborative Research in Computational Neuroscience data sharing website (www.crcns.org) (Figure S3). These recordings are a generous contribution by György Buzsáki’s laboratory (HC3 dataset, Mizuseki et al., 2014).

### Data collection

Recording sessions were performed in an open field (1 m × 1 m) and lasted 4-5 hours. Raw signals were amplified (200x), filtered (1–7.8 kHz) and digitized at 25 kHz (RHA2116, IntanTech). The LFP was obtained by further filtering between 1 – 500 Hz and downsampling to 1000 Hz.

### Data analysis

Active waking and REM sleep periods were identified from spectral content (high theta/delta power ratio) and video recordings (movements during active waking; clear sleep posture and preceding slow-wave sleep for REM). Results were identical for active waking and REM epochs; throughout this work we only show the latter.

We used built-in and custom written MATLAB routines. Filtering was obtained by means of a finite impulse response (FIR) filter using the *eegfilt* function from EEGLAB toolbox (Delorme and Makeig, 2004). The phase time series was obtained through the Hilbert transform. To estimate the instantaneous theta phase of actual data, we filtered the LFP between 4 – 20 Hz, a bandwidth large enough to capture theta wave asymmetry (Belluscio et al., 2012). Similar results were obtained when estimating theta phase by the interpolation method described in Belluscio et al. (2012) (Figure S2). The CSD signals analyzed in Figure S4 were obtained as – A+2B–C for adjacent probe sites. In Figure S5, the independent components were obtained as described in Schomburg et al. (2014); phase-amplitude comodulograms were computed as described in Tort et al. (2010).

### n:m phase-locking

We measured the consistency of the phase difference between accelerated time series (Δ*φ*_*nm*_(*t*_*j*_) = *n* * *φ*_*γ*_(*t*_*j*_)). To that end, we created unitary vectors whose angle is the instantaneous phase difference (*e*^*i*Δ*φ*^_*nm*_^(*t*_*j*_)^), and then computed the length of the mean vector: 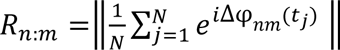. R_n:m_ equals 1 when Δ*φ*_*nm*_ is constant, and 0 when Δ*φ*_*nm*_ is uniformly distributed. This metric is also commonly referred to as “mean resultant length” or “mean radial distance” (Belluscio et al., 2012; Stujenske et al., 2014; Zheng et al., 2016). Qualitatively similar results (i.e., lack of statistically significant n:m phase-locking) were obtained when employing the pairwise phase consistency metric described in Vinck et al. (2010) (not shown). Phase-phase plots were obtained by first binning theta and gamma phases into 120 bins and next constructing 2D histograms of phase counts.

### Surrogates

In all cases, theta phase was kept intact while gamma phase was mocked in three different ways: (1) *Time Shift:* the gamma phase time series is randomly shifted between 1 and 200 ms; (2) *Random Permutation*: a contiguous gamma phase time series of the same length as the original is randomly extracted from the same session. (3) *Phase Scrambling:* the timestamps of the gamma phase time series are randomly shuffled (thus not preserving phase continuity). For each case, R_n:m_ values were computed using either Δ*φ*_*nm*_ distribution for single surrogate run (*Single Run Distribution*) or the pooled distribution of Δ*φ*_*nm*_ over 100 surrogate runs (*Pooled Distribution*).

For each animal, behavioral state (active waking or REM sleep) and epoch length, we computed 300 *Original* R_n:m_ values using different time windows along with 300 mock R_n:m_ values per surrogate method. Therefore, in all figures each boxplot was constructed using the same number of samples (=300 × number of animals). For instance, in Figure 3C we used n=7 animals × 300 samples per animal =2100 samples (but see *Statistics* below). In Figure 2, boxplot distributions for the simulated data were constructed using n=2100.

### Statistics

For simulation results (Figure 2F), given the large sample size (n=2100) and independence among samples, we used one-way ANOVA with Bonferroni posthoc test. For statistical analysis of real data (Figure 3C), we avoided nested design and inflation of power and used the mean R_n:m_ value per animal. In this case, due to the reduced sample size (n=7) and lack of evidence of normal distribution (Shapiro-Wilk normality test), we used the Friedman’s test and Nemenyi post-hoc test.

## Author contributions

A.B.L.T. designed research; R.S.T. acquired and analyzed data; A.B.L.T. and R.S.T. wrote the manuscript.

## Acknowledgment

The authors are grateful to Jurij Brankack and Andreas Draguhn for donation of NeuroNexus probes. Supported by Conselho Nacional de Desenvolvimento Científico e Tecnológico (CNPq) and Coordenaçâo de Aperfeiçoamento de Pessoalde Nível Superior (CAPES). The authors declare no competing financial interests.

## Supplemental Information

**Lack of evidence for cross-frequency phase-phase coupling between theta and gamma oscillations in the hippocampus**

Robson Scheffer-Teixeira & Adriano BL Tort

**Figure S1, Related to Figure 5.**
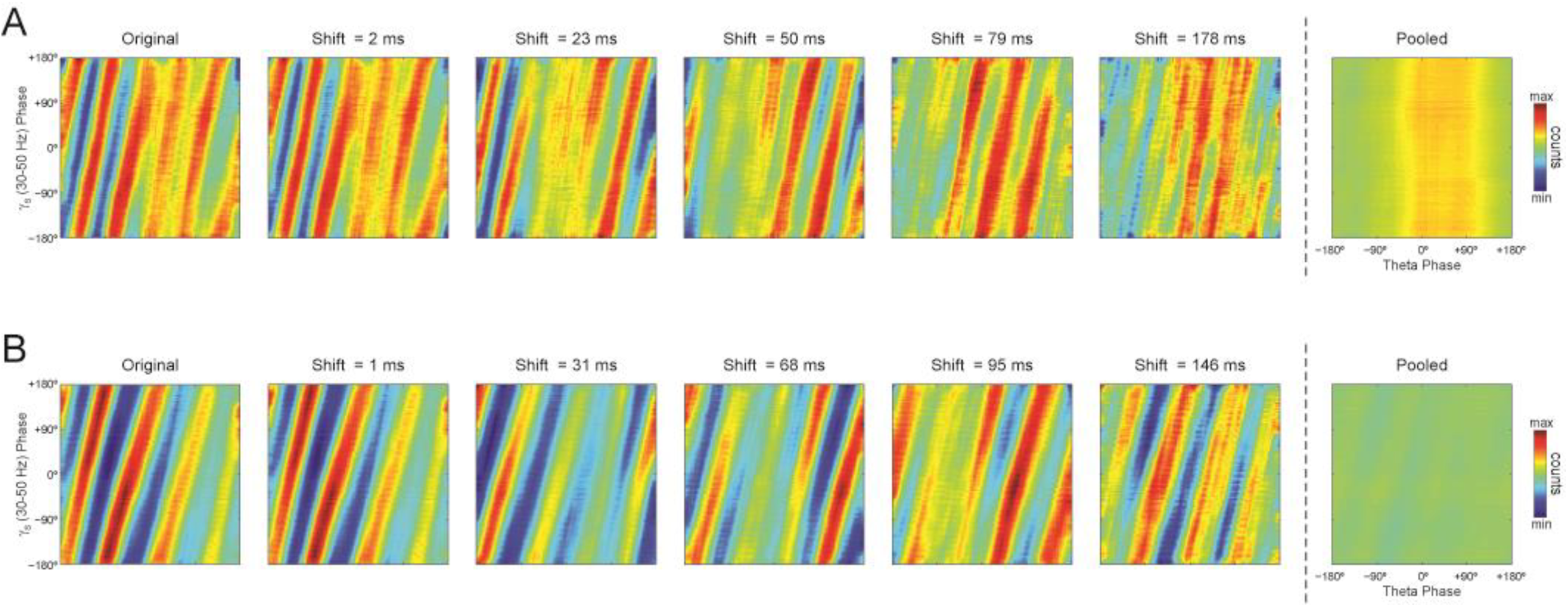
Individual time-shifted surrogate runs exhibit diagonal stripes in phase-phase plots. (A)The middle panels show phase-phase plots for theta and slow gamma computed for different time shifts of the example epoch analyzed in Figure 5A. Notice diagonal stripes in individual surrogate runs. The leftmost panel shows the original phase-phase plot (same as in Figure 5A); the rightmost panel shows the phase-phase plot computed using the pool of all time-shifted surrogate runs (n=1000). (B)Same as above, but for the example epoch analyzed in Figure 5B.

**Figure S2, Related to Figure 3.**
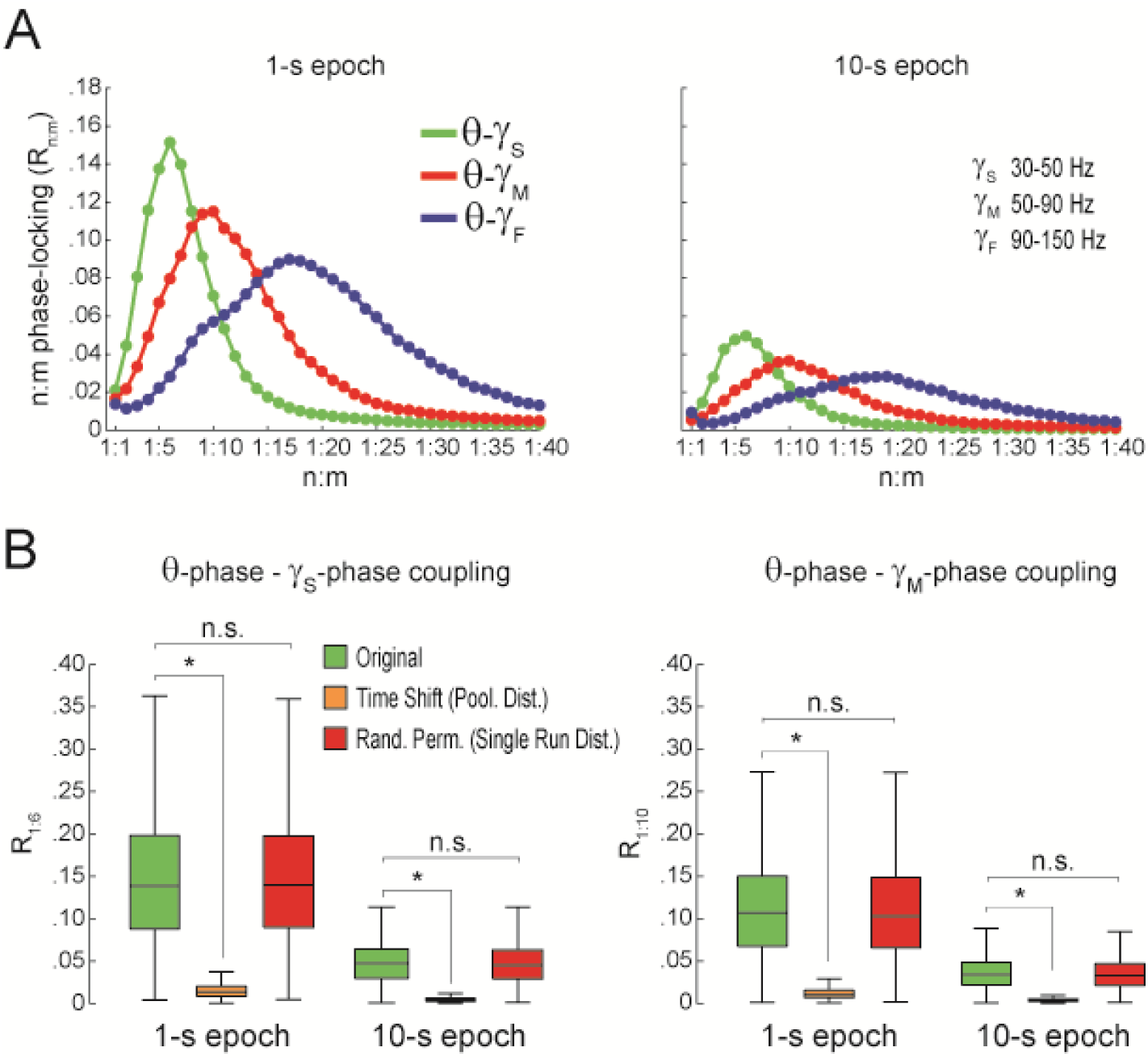
Spurious detection of theta-gamma phase-phase coupling when theta phase is estimated by interpolation. (A)n:m phase-locking levels for actual hippocampal LFPs (same dataset as in Figure 3). Theta phase was estimated by the interpolation method described in Belluscio et al. (2012). (B)Original and surrogate distributions of R_n:m_ values. The original data is significantly higher than surrogate values obtained from pooled Δφ_nm_, but indistinguishable from single run surrogates. ^*^p<0.01, n=7 animals, Friedman’s test with Nemenyi post-hoc test.

**Figure S3, Related to Figure 3.**
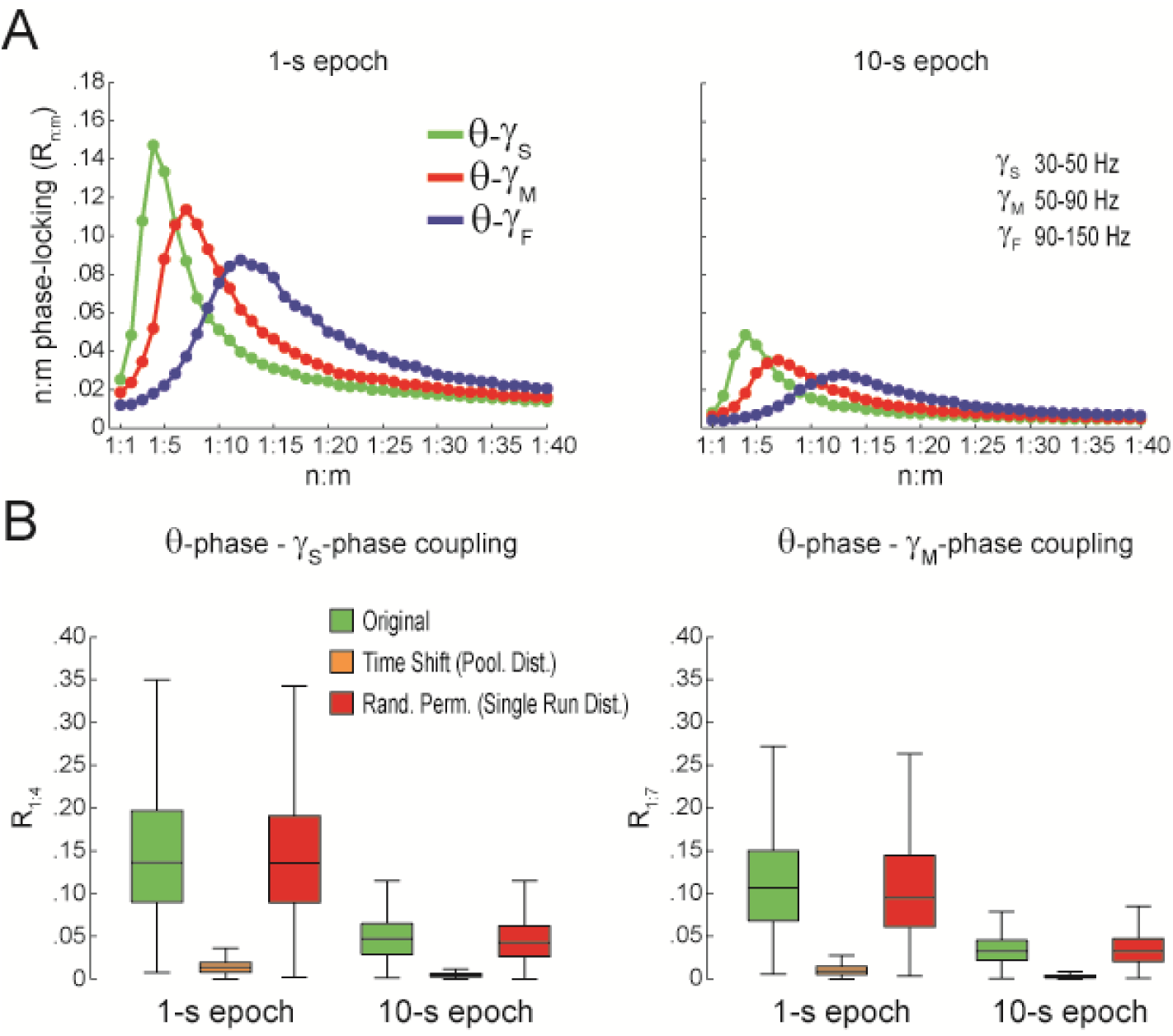
Spurious detection of theta-gamma phase-phase coupling (second dataset). (A)n:m phase-locking levels for actual hippocampal LFPs. (B)Original and surrogate distributions of R_n:m_ values. Results obtained for 3 rats recorded in an independent laboratory (see Material and Methods).

**Figure S4, Related to Figure 3.**
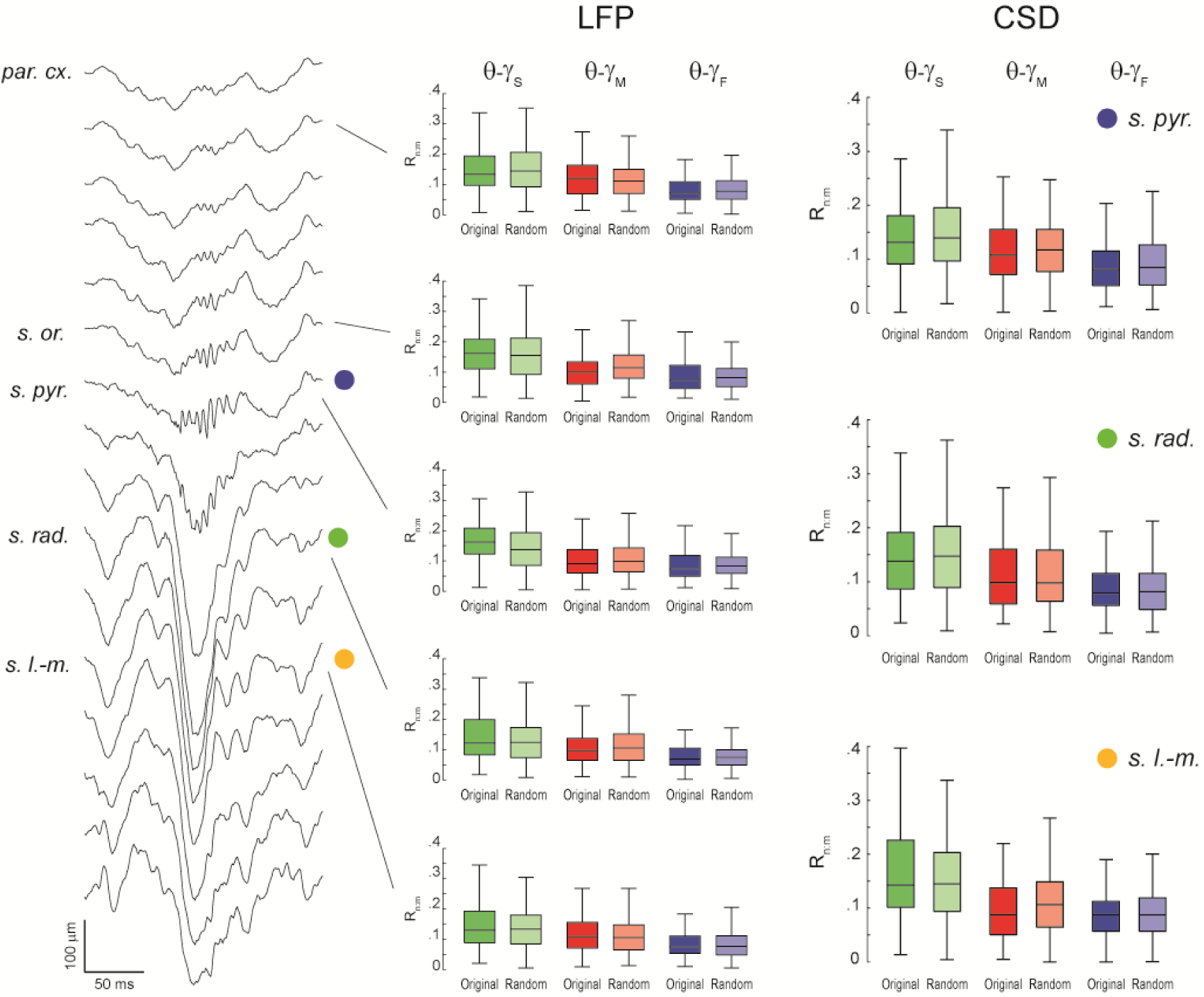
Lack of evidence for theta-gamma phase-phase coupling in all hippocampal layers. (Left) Example estimation of the anatomical location of a 16-channel silicon probe by the characteristic depth profile of sharp-wave ripples (inter-electrode distance = 100 μm). (Middle) Original and surrogate (Random Permutation, Single Run) distributions of R_n:m_ values computed between theta phase and the phase of three gamma sub-bands (1-s long epochs). Different rows show results for different layers. (Right) Distribution of original and surrogate R_n:m_ values computed for current-source density (CSD) signals (1-s long epochs) in three hippocampal layers: s. pyramidale (top), s. radiatum (middle), and s. lacunosum-moleculare (bottom). Notice no difference between original and surrogate values. Similar results were found in all animals.

**Figure S5, Related to Figure 3.**
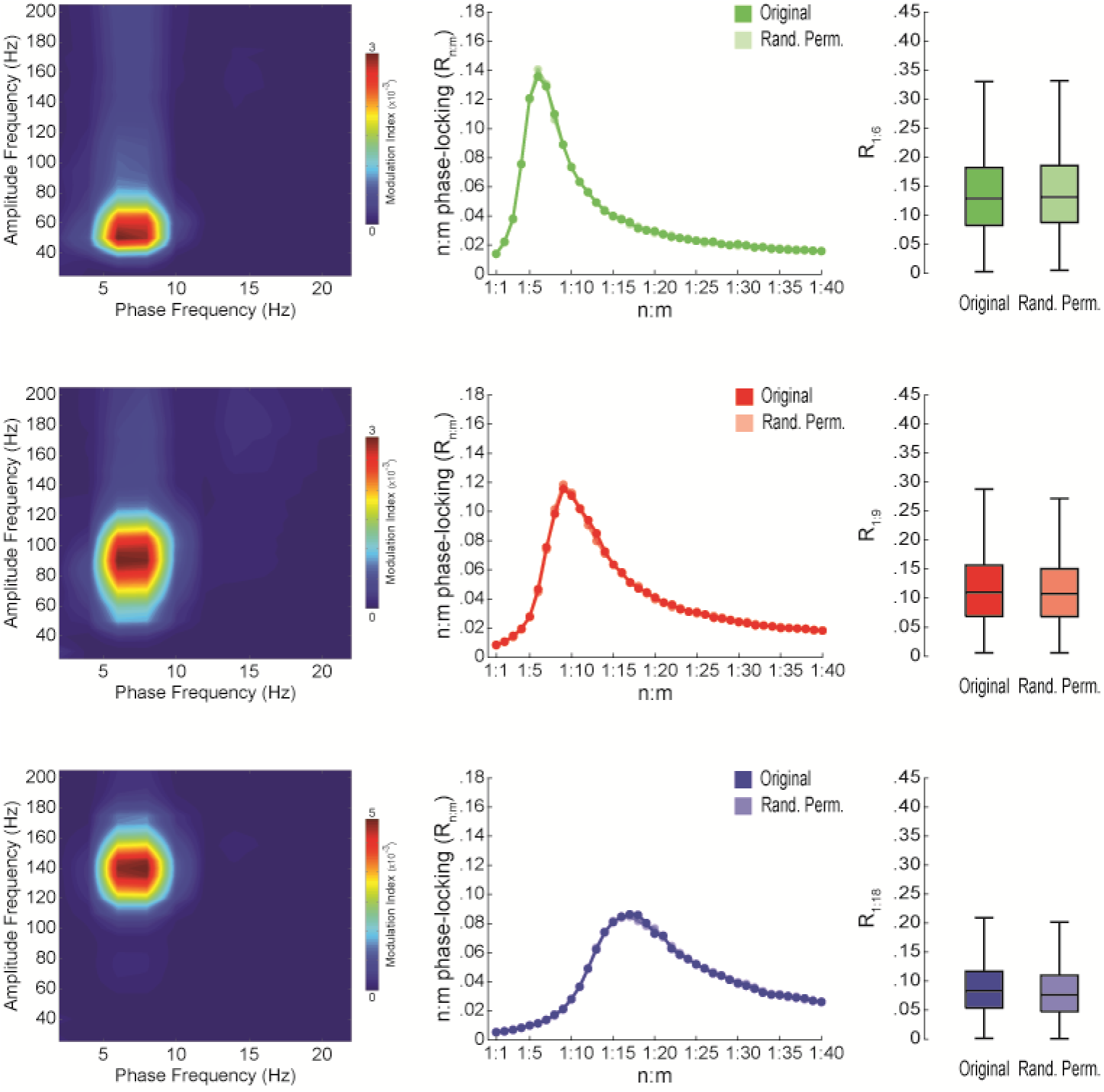
Lack of theta-gamma phase-phase coupling in independent components of gamma activity. (Left) Average phase-amplitude comodulograms for three independent components (IC) that maximize coupling between theta phase and the amplitude of slow gamma (top row), middle gamma (middle row) and fast gamma (bottom row) oscillations (n=4 animals). Each IC is a weighted sum of LFPs recorded in different hippocampal layers (see Schomburg et al., 2014). (Middle) n:m phase-locking levels for theta phase and the phase of ICs filtered at the gamma band maximally coupled to theta in the phase-amplitude comodulogram. (Right) Original and surrogate distributions of R_n:m_ values. R_n:m_ values were computed for 1-s long epochs (n=4 animals); surrogate gamma phases were obtained by Random Permutation/Single Run.

**Figure S6, Related to Figure 3.**
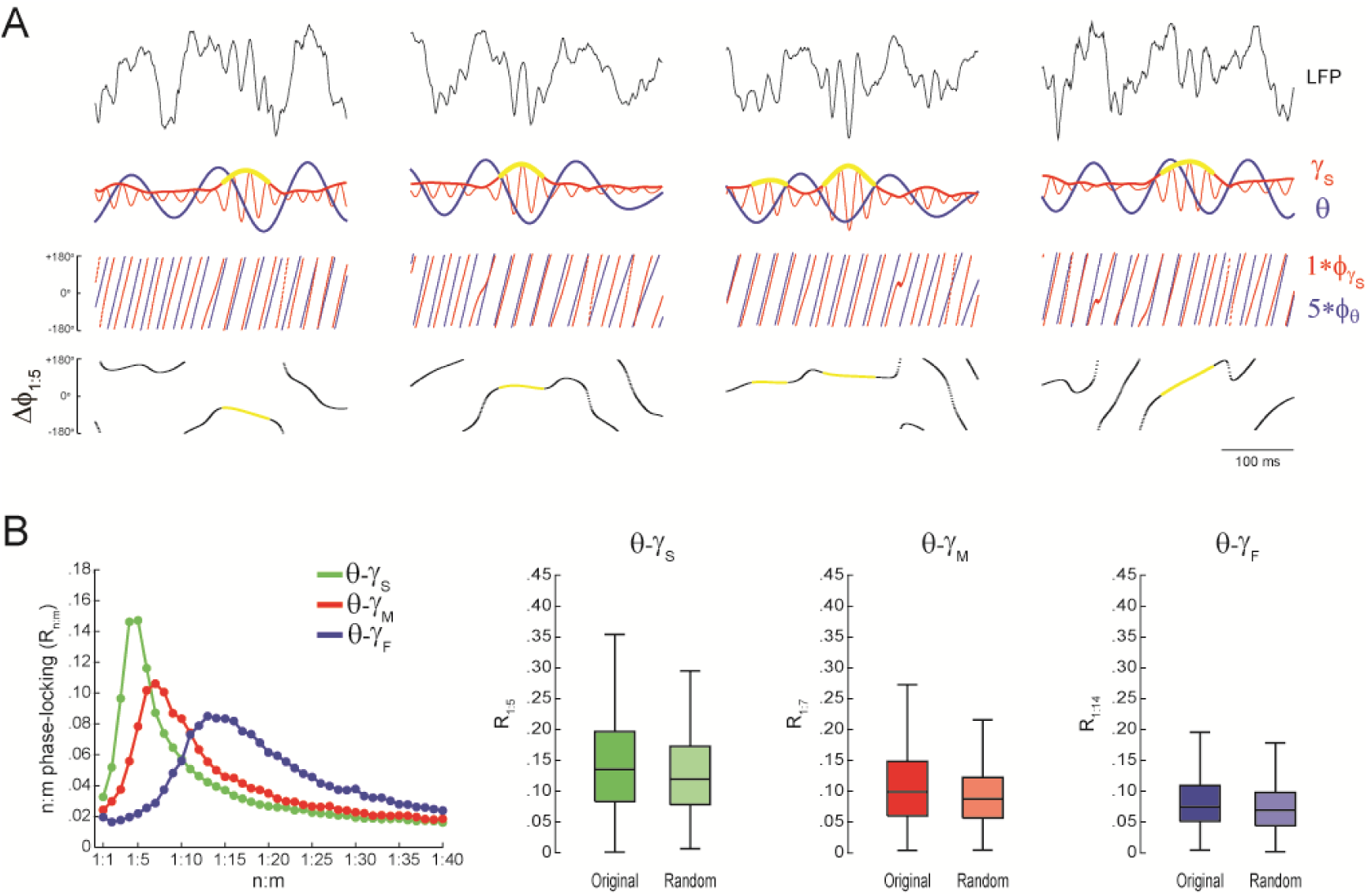
Lack of theta-gamma phase-phase coupling during transient gamma bursts. (A)Examples of slow gamma bursts. Top panels show raw LFPs, along with theta-(thick blue line) and slow gamma-filtered (thin red line) signals. The amplitude envelope of slow gamma is also shown (thick red line). The bottom rows show gamma and accelerated theta phases (m=5), along with their instantaneous phase difference (Δφ_1:5_). For each gamma sub-band, a “gamma burst” was defined to occur when the gamma amplitude envelope was 2SD above the mean. In these examples, periods identified as slow gamma bursts are marked with yellow in the amplitude envelope and phase difference time series. Notice variable Δφ_1:5_ across different burst events. (B)The left panel shows n:m phase-locking levels for theta phase and the phase of different gamma sub-bands (1-s long epochs); for each gamma subband, R_n:m_ values were computed using only theta and gamma phases during periods of gamma bursts. The right panels show original and surrogate (Random Permutation/Single Run) distributions of R_n:m_ values (n=4 animals).

**Figure S7, Related to Figure 4.**
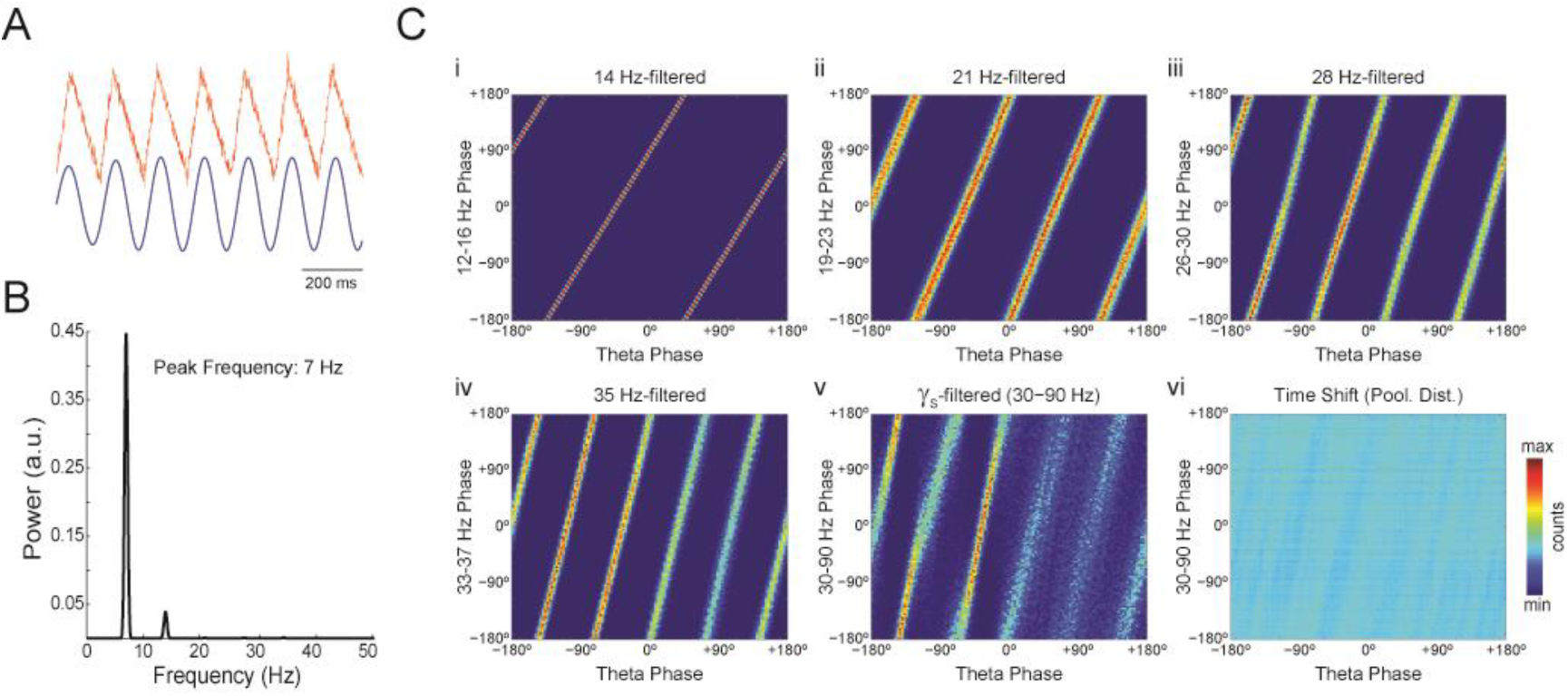
Phase-phase plots of a theta saw-tooth wave with no gamma oscillation display diagonal stripes. (A) Top, LFP simulated as a 7-Hz saw-tooth wave plus noise. No gamma oscillation was added to the signal. Bottom, theta-filtered LFP. (B) Power spectral density. (C) Phasy–phase plots for theta and LFP band-pass filtered at harmonic frequencies (14, 21, 28 and 35 Hz). Also shown are phase-phase plots for the conventional gamma band (30 – 90 Hz) and for pooled surrogate runs.

